# DistilProtBert: A distilled protein language model used to distinguish between real proteins and their randomly shuffled counterparts

**DOI:** 10.1101/2022.05.09.491157

**Authors:** Yaron Geffen, Yanay Ofran, Ron Unger

## Abstract

**Summary:** Recently, Deep Learning models, initially developed in the field of Natural Language Processing (NLP), were applied successfully to analyze protein sequences. A major drawback of these models is their size in terms of the number of parameters needed to be fitted and the amount of computational resources they require. Recently, “distilled” models using the concept of student and teacher networks have been widely used in NLP. Here, we adapted this concept to the problem of protein sequence analysis, by developing DistilProtBert, a distilled version of the successful ProtBert model. Implementing this approach, we reduced the size of the network and the running time by 50%, and the computational resources needed for pretraining by 98% relative to ProtBert model. Using two published tasks, we showed that the performance of the distilled model approaches that of the full model. We next tested the ability of DistilProtBert to distinguish between real and random protein sequences. The task is highly challenging if the composition is maintained on the level of singlet, doublet and triplet amino acids. Indeed, traditional machine learning algorithms have difficulties with this task. Here, we show that DistilProtBert preforms very well on singlet, doublet, and even triplet-shuffled versions of the human proteome, with AUC of 0.92, 0.91, and 0.87 respectively. Finally, we suggest that by examining the small number of false-positive classifications (i.e., shuffled sequences classified as proteins by DistilProtBert) we may be able to identify de-novo potential natural-like proteins based on random shuffling of amino acid sequences.

**Availability:** https://github.com/yarongef/DistilProtBert

## 1 Introduction

In recent years, the field of Natural Language Processing (NLP) has rapidly advanced. Mechanisms and learning paradigms such as attention-based transformers (Vaswani et al., 2017), masked language modeling (Devlin et al., 2018) and special refinements of these methods (Lan et al., 2019; Liu et al., 2019) have improved our understanding of text and our ability to mine it. Transfer learning and pretraining learning procedures using increasingly larger and larger datasets have made it possible to create meaningful representations of words and sentences. Since these techniques often create huge networks, distillation methods have been suggested to create more compact models, while maintaining accuracy (Hinton et al., 2015; Jiao et al., 2019; Sanh et al., 2019). Recently, these NLP techniques have been applied to study protein sequences (Elnaggar et al., 2021; Ofer et al., 2021). Using Recurrent Neural Networks (Alley et al., 2019) and transformer architectures (Brandes et al., 2022; Rives et al., 2021) to create these representations provided approaches to improve the performance in various protein downstream tasks such as secondary structure prediction (Wang Q et al., 2021), flexibility prediction, and fluorescence prediction as shown in TAPE benchmark (Rao et al., 2019). Moreover, recent protein structure prediction algorithms such as AlphaFold (Jumper et al., 2021), RoseTTA fold (Baek et al., 2021) and trRosetta (J. Yang et al., 2020) also utilize deep learning for proteomics, further fueling the rapid development of this field. Here, we show the first application, to the best of our knowledge, of a distilled model for protein sequence analysis. We present DistilProtBert, a novel protein attention-based model based on ProtBert (Elnaggar et al., 2021). By taking advantage of a large pre-training protein dataset, a teacher network, knowledge distillation mechanism and a standard GPU cluster, we demonstrate a simpler yet accurate way to represent proteins. We show that DistilProtBert performs on-par with the full version of ProtBert on two published benchmarks.

In addition, this model may be used to distinguish between real and random proteins. The number of different proteins in nature is huge, estimated to be about 10^11^. Nevertheless, this is only a miniscule fraction of the number of possible sequences that can be created from an alphabet of 20 amino acids, which is 20^N^ where N is the length of the protein, typically several hundred amino acids. The common wisdom is that most of the random sequences will neither fold nor have any useful function. This observation raises an obvious yet fundamental question: What are the characteristics of a sequence of amino acids that result in a protein? In more concrete terms, we asked whether using a machine learning approach, we could distinguish between real sequences of proteins and their shuffled random counterparts. Proteins have characteristic amino acid compositions not only on the single amino acid level, but also at the level of doublets and triplets, and thus, amino acid composition could serve as a key to identifying real proteins (Pe’er et al., 2004). Nevertheless, for a given amino acid composition, distinguishing between real and random sequences that maintain the same amino acid composition remains a major challenge. We show here that DistilProtBert does very well at this challenge and was able to distinguish between real proteins and their shuffled counterparts suggesting that real proteins have underlying characteristics that are not captured by their amino acid composition.

## 2 Methods

### 2.1 Data

#### 2.1.1 Pretraining dataset

ProtBert was pretrained on ~216M proteins from the UniRef100 dataset (Suzek et al., 2007). DistilProtBert was pretrained on ~43M proteins from UniRef50 (Suzek et al., 2007) filtered by length from 20 to 512 amino acids.

#### 2.1.2 Benchmarks datasets

DistilProtBert was evaluated on several benchmark tasks. For secondary structure prediction (Q3), we used Netsurfp2 (Klausen et al., 2019), CASP12 (Moult et al., 2018), CB513 (Cuff & Barton, 1999) and TS115 (Y. Yang et al., 2016) datasets, and for membrane-bound vs water soluble prediction (Q2) we used the DeepLoc dataset (Almagro Armenteros et al., 2017). To distinguish real protein sequences from random k-let shuffled versions of them, we used singlet, doublet, and triplet non-redundant versions of the human proteome.

### 2.2 Pretraining

DistilProtBert was pretrained on ~43M sequences from UniRef50 with length ranging from 20 to 512 amino acids. We pretrained the model on a masked language modeling task with masking probability of 15%. Pretraining was done on five v100 32GB Nvidia GPUs from a DGX cluster with a local batch size of 16 examples. We used AdamW optimizer with an initial learning rate of 5e^-5^ and no weight decay. The model was trained for 3 epochs using mixed precision, and dynamic padding. Every epoch run took approximately 4 days, resulting in total pretraining time of 12 days.

### 2.3 Knowledge distillation

The weights of DistilProtBert were initialized using the weights learned from ProtBert. Knowledge from the teacher network (ProtBert), was distributed towards the student network (DistilProtBert), via the loss function. As described in (Sanh et al., 2019) the loss is comprised of 3 equally weighted parts: *L_mlm_, L_ce_* and *L_cos_*. A softmax temperature of 2 was set during pretraining (Hinton et al., 2015).

### 2.4 Fine tuning for benchmark tasks

To fine-tune DistilProtBert for secondary structure prediction (Q3) an all-tokens (amino acid) classification head (linear layer for each hidden-state output) was added on top of our pretrained model. For membranebound vs. water soluble prediction (Q2), a first token classification head was added on top of our pretrained model. In this strategy, we utilized the first token from DistilProtBert corresponding to [CLS] token, which attends all the other tokens in the sequence to capture the best representation for the whole sequence. Fine-tuning of the model for all benchmark tasks was performed without any layer freezing and was done with the same hyperparameters as reported by (Elnaggar et al., 2021).

### 2.5 Real versus shuffled protein classification task

The real versus shuffled proteins task was constructed in the following manner: singlet, doublet and triplet versions of the human proteome were used as our dataset. Out of 20,577 human proteins (from UniProt), sequences shorter than 20 amino acids or longer than 512 amino acids were removed, resulting in a set of 12,703 proteins. The uShuffle algorithm (Jiang et al.,2008) was then used to shuffle these protein sequences while maintaining their k-let distribution for k=1,2,3. The very few sequences for which uShuffle failed to create a shuffled version were eliminated.

A notorious problem in classification tasks of proteins is that many proteins are part of a family with several homologs and tend to be similar to each other, thereby rendering the distinction between training and test sets meaningless. To address this problem, we ran all the sequences (real and shuffled) through the h-CD-HIT algorithm, which is the most rigorous version of CD-HIT. Three subsequent filter stages of CD-HIT were performed with pairwise identity cutoffs of 0.9, 0.5 and 0.1 respectively. Note that this filtering resulted in a much smaller number of sequences that maintained triplet frequency, as it was apparently difficult to shuffle sequences to deviate sufficiently from their source while maintaining the triplet composition. All the sets contained pairs of sequences, the real protein and its shuffled counterpart, and the algorithm was tasked with classifying the real versus the random sequence. We split the sequences to training and test sets, 80% were set as a training set, and the rest 20% were set aside for test. A 10-fold cross validation was performed for each one of the k-let training sets and the average results were reported. Then, the performance of our model was evaluated on the 20% test set that was set aside. The datasets sizes are shown in Table 1.

**Table 1.** Datasets sizes after each filtering stage

| k-let | h-CD-HIT | Training set | Validation set | Test set |
| --- | --- | --- | --- | --- |
| Singlet | 11,698 | 8,424 | 936 | 2,338 |
| Doublet | 11,658 | 8,388 | 932 | 2,338 |
| Triplet | 3,688 | 2,664 | 296 | 728 |

DistilProtBert was used as a feature extractor, performing max pooling (filter size=16, stride=16) on each of the token (amino acid) representations. Afterwards, concatenation and zero padding was performed. We then used a feed-forward neural network to classify the sequences into the two classes. We trained the network with the following hyperparameters: batch size 32, RAdam optimizer, learning rate of 0.000001, dropout 0.1 and 100 epochs with early stopping.

## 3 Results

The general concept of knowledge distillation is to teach a smaller network, dubbed the student network, to mimic a larger network, dubbed the teacher network. By training a student network on a masked language modeling task (MLM) and narrowing the gap to the teacher network parameter distribution, it is possible to create a distilled network that maintains a high level of performance. Bert-based models, such as ProtBert, are amenable to this procedure, since they are very large and resource consuming in terms of hardware and running time. Therefore, to efficiently utilize such models, distilled versions are commonly developed (Jiao et al., 2019; Sanh et al., 2019; Sun et al., 2020).

### 3.1 Optimization of the model and memory consumption

Inspired by DistilBert that was designed for natural language processing tasks (Sanh et al., 2019) and using ProtBert (Elnaggar et al., 2021) as a teacher network, we created a distilled ProtBert implementation which we termed “DistilProtBert”. While ProtBert contains 420M parameters, we were able to reduce the number of DistilProtBert parameters by almost half, to 230M. In addition, we were able to pretrain DistilProtBert on commodity hardware of five v100 Nvidia 32GB GPUs from a single DGX cluster. This is a 51-fold improvement from ProtBert pretraining, which required a TPU Pod of 64 nodes with 512 16GB TPUs in total. Consequently, fine-tuning and inference times on downstream tasks were twice as fast as those attained using ProtBert. Moreover, in DistilProtBert only short sequences of 512 amino acids in length were used during pretraining, as opposed sequences of up to 2048 amino acids in length that were used for ProtBert pretraining.

### 3.2 Deep learning architecture

The student network, DistilProtBert, and teacher network, ProtBert, share the same general architecture. They both have 1024 neurons in each hidden layer, 4096 neurons in each intermediate hidden layer, and 16 attention heads in each one of their hidden layers. Yet, the number of total hidden layers in DistilProtBert was smaller; in contrast to 30 hidden layers in ProtBert, only 15 hidden layers were used, taking each layer alternately.

### 3.3 Evaluation tasks

We first evaluated DistilProtBert performance on two tasks that were previously studied using ProtBert (Elnaggar et al., 2021), secondary structure prediction (Q3), and prediction of membrane-bound vs water soluble proteins (Q2). The results are shown in Table 2.

**Table 2.**
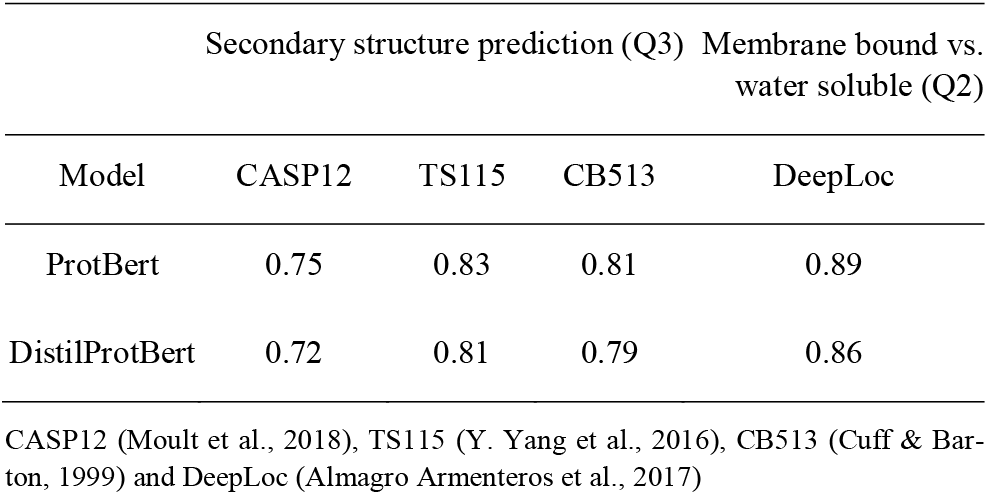
Accuracy results for benchmark tasks

For both tasks, DistilProtBert demonstrated results on-par with ProtBert-UniRef100, with a maximal reduction of only 3 points. Yet, training and inference times were twice as fast when using the same hyperparameters as in ProtBert’s fine-tuning. Further fine-tuning may improve results.

We then turned to the challenge of distinguishing real proteins from their shuffled counterparts. We first tried to address this challenge by using a bidirectional LSTM network. This model achieved poor results of 0.71, 0.68 and 0.61 AUC for the singlet, doublet and triplet human proteome test sets respectively. We then investigated the performance of DistilProtBert on this task. Use of this model resulted in dramatic improvements (Tables 3 and 4), demonstrating that 1) deep learning models can perform remarkably well in this difficult task, and 2) the performance of our distilled version approaches that of the full ProtBert version. Note that the results are very good both on the cross validation and on the test sets.

**Table 3.** Cross validation scores

| Dataset | Model | Accuracy | F1 | Precision | Recall | AUC |
| --- | --- | --- | --- | --- | --- | --- |
| Singlet | LSTM | 0.71 | 0.65 | 0.82 | 0.54 | 0.71 |
|  | ProtBert | 0.92 | 0.92 | 0.97 | 0.88 | 0.92 |
|  | DPB | 0.91 | 0.91 | 0.95 | 0.87 | 0.91 |
| Doublet | LSTM | 0.69 | 0.63 | 0.77 | 0.54 | 0.69 |
|  | ProtBert | 0.91 | 0.90 | 0.96 | 0.85 | 0.91 |
|  | DPB | 0.89 | 0.89 | 0.93 | 0.85 | 0.89 |
| Triplet | LSTM | 0.59 | 0.57 | 0.61 | 0.56 | 0.59 |
|  | ProtBert | 0.93 | 0.93 | 0.97 | 0.89 | 0.93 |
|  | DPB | 0.87 | 0.86 | 0.91 | 0.82 | 0.87 |

DPB = DistilProtBert

**Table 4.** Test scores

| Dataset | Model | Accuracy | F1 | Precision | Recall | AUC |
| --- | --- | --- | --- | --- | --- | --- |
| Singlet | LSTM | 0.71 | 0.66 | 0.80 | 0.56 | 0.71 |
|  | ProtBert | 0.93 | 0.93 | 0.96 | 0.89 | 0.93 |
|  | DPB | 0.92 | 0.92 | 0.94 | 0.90 | 0.92 |
| Doublet | LSTM | 0.68 | 0.62 | 0.78 | 0.51 | 0.68 |
|  | ProtBert | 0.92 | 0.92 | 0.97 | 0.87 | 0.92 |
|  | DPB | 0.91 | 0.90 | 0.94 | 0.87 | 0.91 |
| Triplet | LSTM | 0.61 | 0.57 | 0.64 | 0.51 | 0.61 |
|  | ProtBert | 0.92 | 0.92 | 0.98 | 0.87 | 0.92 |
|  | DPB | 0.87 | 0.86 | 0.89 | 0.84 | 0.87 |

DPB = DistilProtBert

## 4 Discussion

In recent years, it was shown that the use of pretrained transformers for protein language-based prediction models is very effective. In accordance with this trend, we developed DistilProtBert, a novel protein language model. This model provided us with a more efficient way to create concise protein representations, while maintaining good performance, and allowed us to pretrain and fine-tune a protein language model without the need for expensive high-performance computing systems. Thus, it provides an accessible and more convenient means to create protein representations.

As we described above, DistilProtBert can be used on commodity hardware to perform several downstream tasks, including secondary structure prediction and the classification task distinguishing real versus shuffled amino acid protein sequences. It enables fine-tuning and feature extraction in a fast and efficient manner. As access to such models becomes easier (Wolf et al., 2019) other BERT-like architectures, for example TinyBert (Jiao et al., 2019) and MobileBert (Sun et al., 2020), should be examined.

In addition, we performed a novel classification task between real proteins and their k-let shuffled versions. The success of the attention based models in this task suggests that there is a hidden structure in real proteins that is not reflected in their k-let amino acid composition. We showed that standard neural network architectures, such as LSTM, achieved much lower performance and thus it seems that attention-based models can capture properties of proteins that are not found by classical machine learning schemes. In addition to the technical success, this achievement can teach us that there are characteristics of real proteins that are not reflected in the k-let amino acid composition. This is a profound statement since other properties that define proteins such as secondary structure preference are affected, to a large extent, by the local amino acid composition. Since our “words” are preserved on the triplet amino acid level, we may conclude that proteins have long range dependencies between amino acids that DistilProtBert was able to detect. Elucidating and understanding these underlying dependencies from the model would not be a simple endeavor as deep learning schemes are notoriously difficult to interpret. However, there are efforts, for example (Vig et al., 2020), that may be able to facilitate gaining biological knowledge from deep learning schemes.

Even without fully understanding the defining characteristics of valid proteins, our model can be used to identify good starting points for de-novo protein design. This can be done by examining the small number of false-positive classifications i.e., the shuffled amino acid sequences that were classified as natural proteins by DistilProtBert. It would be interesting to determine, by synthesizing several of these proteins, if indeed they exhibit protein-like properties in terms of folding and stability.

## Acknowledgements

We thank the Data Science Institute (DSI) of Bar-Ilan University for a research grant and use of computer resources.

